# Zinc can counteract selection for ciprofloxacin resistance

**DOI:** 10.1101/780981

**Authors:** Michiel Vos, Louise Sibleyras, Lai Ka Lo, Elze Hesse, William Gaze, Uli Klümper

**Author notes:** corresponding author: Uli Klümper, CLES & ESI University of Exeter, TR109FE Penryn, United Kingdom.

## Abstract

Antimicrobial resistance (AMR) has emerged as one of the most pressing global threats to public health. AMR evolution occurs in the clinic but also in the environment, where low concentrations of antibiotics and heavy metals can respectively select and co-select for resistance. While the selective potential for AMR of both antibiotics and metals is increasingly well-characterized, studies exploring the combined effect of both types of selective agents are rare. It has previously been demonstrated that fluoroquinolone antibiotics such as ciprofloxacin can chelate metal ions. To investigate how ciprofloxacin resistance is affected by the presence of metals, we quantified selection dynamics between a ciprofloxacin-susceptible and an isogenic ciprofloxacin-resistant *Escherichia coli* MG1655 strain across a gradient of ciprofloxacin concentrations in the presence and absence of Zinc cations (Zn^2+^). The minimal selective concentration (MSC) for ciprofloxacin resistance significantly increased up to 5-fold in the presence of Zn^2+^. No such effect on the MSC was found for gentamicin, an antibiotic not known to chelate zinc cations. Environmental pollution usually consists of complex mixtures of antimicrobial agents. Our findings highlight the importance of taking antagonistic as well as additive or synergistic interactions between different chemical compounds into account when considering their effect on bacterial resistance evolution.

**Graphical abstract:** 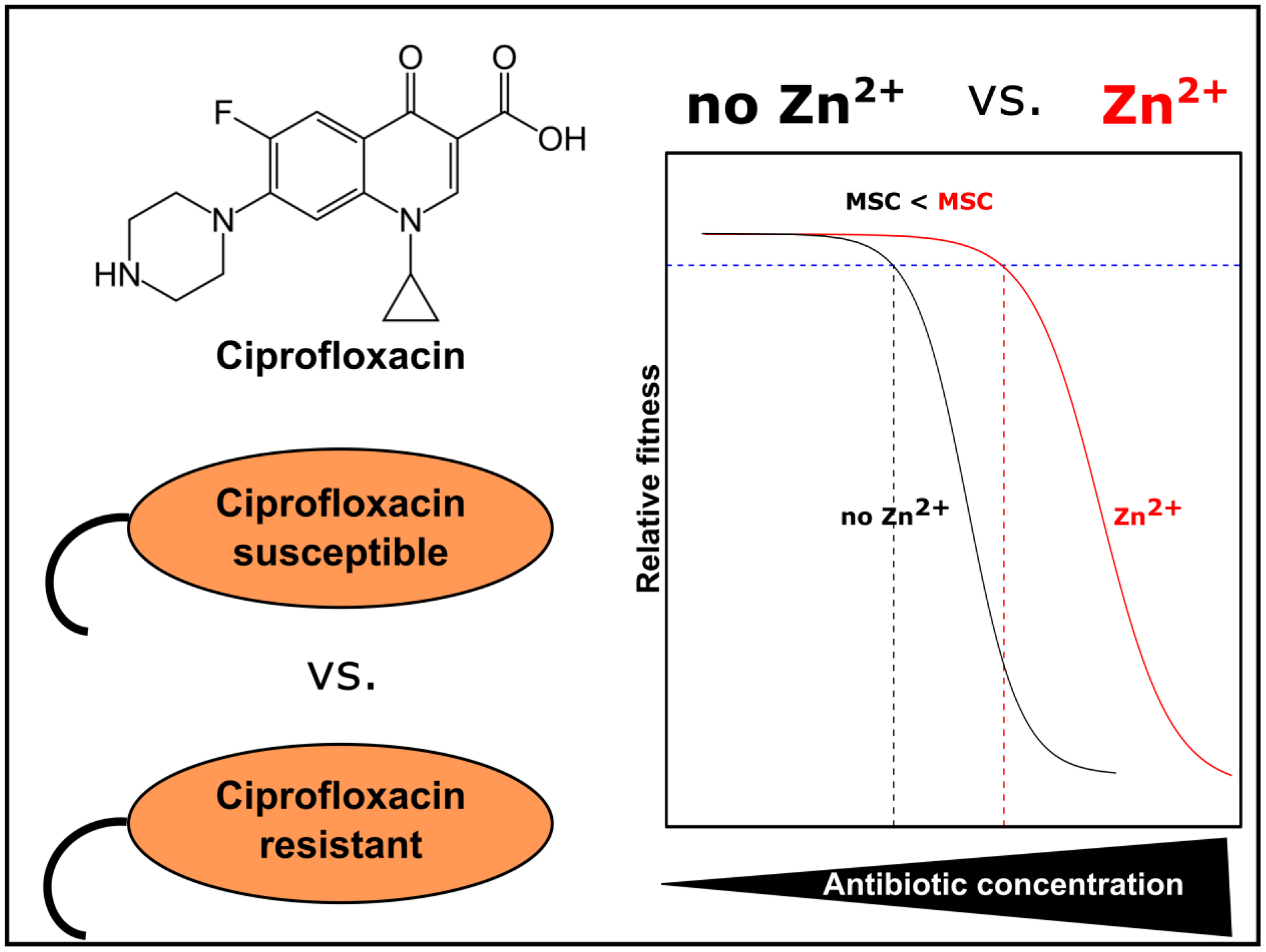

**One sentence summary:** The minimal selective concentration for a ciprofloxacin resistant *E. coli* strain increases up to 5-fold in the presence of Zinc cations.

## Introduction

The emergence and spread of antimicrobial resistance (AMR) genes in bacterial pathogens constitutes a major threat to human health (WHO 2014). Although AMR genes are ancient and have evolved as a result of microbial interactions, as for example evidenced by their presence in permafrost samples minimally impacted by anthropogenic activity (D’Costa *et al*. 2011), the misuse of antibiotics in healthcare and agriculture has caused a rapid and worrying increase in their prevalence and mobility (Knapp *et al*. 2010). The environmental dimensions of AMR evolution are increasingly appreciated (Wellington *et al*. 2013; Larsson *et al*. 2018; Smalla *et al*. 2018), with two major research strands emerging. First, recent studies utilising both single species (Gullberg *et al*. 2011, 2014; Liu *et al*. 2011; Klümper *et al*. 2019b) and complex microbial communities (Lundström *et al*. 2016; Murray *et al*. 2018) have demonstrated that selection for AMR can occur at antibiotic concentrations much lower than those that prevent the growth of susceptible bacteria. These studies highlight the importance of considering the Minimum Selective Concentration (MSC) as opposed to the Minimum Inhibitory Concentration (MIC) for assessing risks associated with antibiotic concentrations. Second, environmental pollution with non-antibiotic compounds such as biocides or metals can contribute to the spread and selection of AMR through processes such as cross-resistance, co-selection and co-regulation (Baker-Austin *et al*. 2006; Pal *et al*. 2017; Dickinson *et al*. 2019) or by altering transfer dynamics of AMR plasmids (Klümper *et al*. 2017, 2019a). Pollution with metals is especially problematic as metals are highly persistent and toxic even at low concentrations (Gadd and Griffiths 1977). In certain environmental settings, heavy metals such as Cu and Zn may even constitute stronger selective agents for antibiotic resistance than the antibiotic itself (Song *et al*. 2017).

While the selective potential for AMR of antibiotics and heavy metals has been relatively well-characterized, experimental studies exploring their combined effect on resistance evolution are rare. The presence of a second antibiotic could, for example, either potentiate or decrease antibiotic efficacy (Cao *et al*. 2012; Churski *et al*. 2012) and hence cause a decrease or increase in MSC. Metals and antibiotics can similarly have synergistic or antagonistic effects. For example, metal complexation can decrease the hydrolysis potential of β-lactam antibiotics by β-lactamases and hence increases antibiotic potency (Anacona 2001). In contrast, the selective potential of more general mechanisms conferring simultaneous resistance to metals and antibiotics, such as efflux pumps, can be increased (Mata, Baquero and Pérez-Díaz 2000; Aendekerk *et al*. 2002). It is also possible that metals could diminish the activity of antibiotics by binding and inactivation (Li, Nix and Schentag 1994). As metal-antibiotic interactions are varied and highly relevant in pollution scenarios, it is important to address how antibiotics and metals jointly affect bacterial resistance spread.

Fluoroquinolones are recognized as critically important antibiotics for human health by the WHO (WHO 2016) and are characterized by a high degree of persistence in the environment (Kümmerer, Al-Ahmad and Mersch-Sundermann 2000). Concentrations in environmental settings range from low ng/L to μg/L, while exceptionally high levels of ciprofloxacin in the mg/L range have been found in effluents from drug manufacturers and in nearby industrially polluted environments (Gothwal and Shashidhar 2015). Interactions between fluoroquinolones and metals have received previous attention but with mixed results. Metal(II)-ciprofloxacin complexes were shown to have better solubility and greater activity against gram-positive and gram-negative pathogens compared to uncomplexed ciprofloxacin (Chohan, Supuran and Scozzafava 2005; Imran *et al*. 2007; Patel, Chhasatia and Parmar 2010). However, a host of studies has also demonstrated that metal-chelated ciprofloxacin, even whilst more readily transported across the bacterial cell membrane, has reduced antimicrobial efficacy (Li, Nix and Schentag 1994; Ma, Chiu and Li 1997; Seedher and Agarwal 2010).

To test whether metal chelation could impact selection for antibiotic resistance, we here quantified growth rates of a ciprofloxacin-susceptible and an isogenic ciprofloxacin-resistant *Escherichia coli* MG1655 strain across a gradient of ciprofloxacin concentrations in the presence and absence of Zinc cations (Zn^2+^). We also performed a competition experiment between both strains in mine-waste contaminated stream water to test whether complex environmental mixtures of metals could affect selection for ciprofloxacin resistance. Our results shed light on a hitherto unappreciated dimension of environmental AMR selection, namely that there is a selective window where metal pollution may reduce the selective effects of antibiotic pollution.

## Material and Methods

### Strains

The ancestral, and antibiotic susceptible *Escherichia coli* MG1655 strain was chromosomally tagged with a Tn*7* gene cassette encoding constitutive red fluorescence, expressed by the *mCherry* gene (Klümper, Dechesne and Smets 2014; Klümper *et al*. 2015). Isogenic pairs of this strain and its resistant counterpart were created for two antibiotics, ciprofloxacin and gentamicin. The ciprofloxacin resistant strain was created through spontaneous chromosomal mutation by evolving the susceptible strain by serial inoculation of overnight culture in LB medium supplemented with incremental concentrations of ciprofloxacin (0.0625, 0.125, 0.25, 0.5 and 1 µg/mL), until a resistant phenotype evolved able to grow at the highest concentration. A gentamicin resistant variant of the *E. coli* ancestral strain was tagged through electroporation with the pBAM delivery plasmid containing the mini-Tn*5* delivery system (Martínez-García *et al*. 2011) for gentamicin resistance gene *aacC1* encoding a gentamicin 3′-*N*-acetyltransferase (Kovach *et al*. 1995). The ciprofloxacin- and gentamicin-resistant strains as well as the ancestral susceptible strain were grown overnight in sterile LB broth supplemented with the appropriate antibiotic (0.5 μg/mL ciprofloxacin, 10 μg/mL gentamicin), harvested by centrifugation (3,500xg, 20min, 4°C), three times washed to remove residual antibiotics with the finally density adjusted to OD_600_ 0.1 (∼10^7^ bacteria/mL) in sterile 0.9% NaCl solution for use in experiments.

### Growth rate assays

Individual strains were inoculated at ∼10^5^ cells/mL in triplicate into LB broth supplemented with ZnSO_4_ (0.0 mM, 0.5 mM, 1 mM) and either ciprofloxacin (eleven two-fold decreasing concentrations starting at 0.4 μg/mL) or gentamicin (eleven two-fold decreasing concentrations starting at 10 μg/mL). After 24h growth under shaking conditions (120 rpm) OD_600_ was measured to determine bacterial growth using a Synergy 2 spectrophotometer (Biotek Instruments Inc., VT, USA). Individual growth rates of the strains were calculated relative to growth of the susceptible strain in a control treatment unamended with zinc or antibiotics.

Individual fitness values (*ρ*) at each concentration were calculated using the growth rate (*γ*) of the susceptible (s) and the resistant (r) strain at each individual concentration, with bacterial numbers (n) calculated based on OD_600_ measurements.

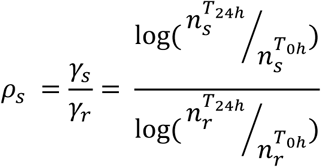

Statistical significance testing (n=3) was performed using a two-tailed *t*-test to test whether for each concentration relative fitness of the sensitive strain differs from a scenario with no selection (*ρ*=1) or if relative growth significantly differed from that of the susceptible strain in the absence of antibiotics and zinc. To compare the average relative fitness of different samples an unpaired two sample *t*-test was performed.

The experimental data linking fitness to antibiotic exposure at the different metal doses were fitted with a four parameter log-logistic dose-response curve using the ‘drc’ (Analysis of Dose-Response Curves) package in R (Knezevic, Streibig and Ritz 2007) with separate curves fitted for each level of zinc, using ‘curveid’ with Zn^2+^ as factor. To integrate the no antibiotic control (c = 0 μg/mL) in the log-logistic model its concentration was set to a minimum threshold as per the developers’ instructions (i.e. by dividing lowest non-zero value by 10). Using these modelled curves the minimal selective concentration at which *ρ*_s_ = *ρ*_r_ = 1 was calculated for each scenario, as well as their standard deviation and 95% confidence intervals. To test if Zn^2+^ alters the dose response relationship, the fitted model was compared to a reduced model where fitness data were pooled across Zn^2+^ concentrations.

### Competition assay

The gentamicin-resistant, ciprofloxacin-susceptible strain was competed with the isogenic gentamycin-sensitive, ciprofloxacin-resistant strain. The pair was inoculated at a 1:1 ratio and initial density of ∼10^5^ cells/mL into 10 mL of LB broth supplemented with ZnSO_4_ (0.0 mM, 0.5 mM) and ciprofloxacin (0.001 μg/mL, 0.01 μg/mL, 0.025 μg/mL, 0.1 μg/mL) in triplicate 30mL glass vial microcosms. Microcosms were incubated shaken (120 rpm) at 37°C for 24h, which allowed growth up to carrying capacity. 100 μL (1%) was then transferred daily for seven days to a fresh microcosm. From each reactor after 7 days (T_7d_), as well as the inoculum (T_0d_), a dilution series in sterile 0.9% NaCl solution was prepared and plated on LB containing either 10 μg/mL gentamicin (selective for ciprofloxacin-susceptible) or 0.5 μg/mL ciprofloxacin (selective for ciprofloxacin-resistant). Plating of the respective strains on the counter selecting plates did not lead to any growth of spontaneous mutants. The relative fitness of the susceptible strain compared to the resistant strain was calculated based on their individual growth rate throughout the competition experiment taking into account a 10^14^ dilution factor throughout the 7 days of growth. Statistical significance testing (n=3) was performed using a two-tailed *t*-test against 1, to test whether for each concentration relative fitness of the sensitive strain differs from a scenario with no selection (*ρ*=1). To compare the average relative fitness of different strains at each concentration an unpaired two sample *t*-test was performed.

### Water sampling

A water sample was collected in May 2019 from a tributary to the Carnon river near Bissoe, Cornwall, UK (50°13’54.6”N, 5°07’48.7”W). A sterile 1L Duran bottle was filled and immediately transported back to the laboratory to use for the growth rate assay or competition assay on the same day of sampling. The water was supplemented with sterile LB broth powder and sterilized through 0.2 μm^2^ pore filters (Whatman). Sterility was confirmed by incubating 10mL of the liquid medium at 37°C overnight. An aliquot of the water sample was frozen at -20°C and analysed in triplicate for metal content using ICP-MS on an Agilent 7700x (Agilent Technologies, Santa Clara, Ca, USA) at the Camborne School of Mines laboratory at the University of Exeter.

## Results

### Effect of Zinc on selection for ciprofloxacin resistance

To determine the effect of zinc cations on selection for ciprofloxacin resistance, isogenic ciprofloxacin-resistant and ciprofloxacin-susceptible *E. coli* strains were grown in isolation across a gradient of ciprofloxacin and Zn^2+^ concentrations for 24h. The growth rate of each strain was plotted relative to that of the ciprofloxacin-susceptible strain grown in control LB (without ciprofloxacin and Zn^2+^) (Figure 1). A small but significant fitness cost of ciprofloxacin resistance in the absence of antibiotics and metals was apparent (82.8 ± 3.4% relative growth, mean ± SD) (p=0.012, *t*-test against 1) (Figure 1A). This cost of resistance in the absence of ciprofloxacin was also apparent across both Zn^2+^ concentrations tested (0.5 mM: 90.7 ± 5.3%, p=0.093: 1 mM: 83.8 ± 3.7%, p=0.0161, two-tailed *t*-test against 1) (Figure 1B,C). Growth of both strains significantly decreased with increasing Zn^2+^ concentrations in the absence of ciprofloxacin to 78.7 ± 4.1% (0.5 mM, p<0.0001) and 55.4 ± 2.4% (1 mM, p<0.0001).

**Fig. 1:**
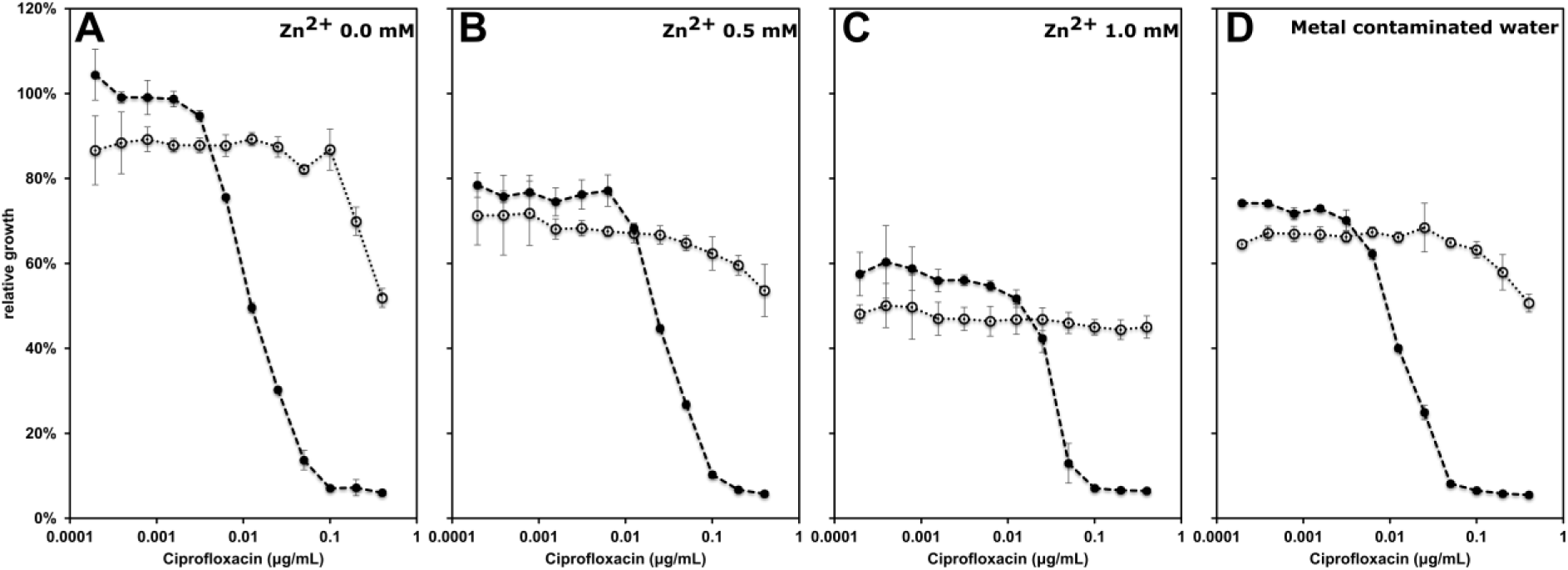
Growth rate of a ciprofloxacin-susceptible strain (filled circles) and a ciprofloxacin-resistant strain (open circles) in LB medium amended with ciprofloxacin and zinc as well as in metal contaminated water supplemented with LB, relative to the growth rate of the ciprofloxacin-susceptible strain in control LB. Growth was quantified based on OD_600_ after 24 h of growth at 37°C (n=3, average ± SD).

A significant growth rate advantage for the ciprofloxacin-resistant strain was detected at 0.00625 μg/mL ciprofloxacin (*ρ*_s,0mM_=0.949 ± 0.001, p=0.0002, two-tailed *t*-test against 1) in the absence of Zn^2+^ (Figure 2). However, at the same antibiotic concentration the growth rate of the ciprofloxacin-resistant strain was lower than the susceptible strain in the presence of 0.5 mM (*ρ*_s,0.5mM_=1.050 ± 0.018; p=0.0412) and lower still when increasing the Zn^2+^ concentration to 1.0 mM (*ρ*_s,1.0mM_=1.073 ± 0.010; p= 0.0059). In the presence of Zn^2+^ the lowest concentration with a significant growth rate advantage for the ciprofloxacin-resistant strain increased from 0.00625 μg/mL ciprofloxacin (no Zn2+) to 0.025 μg/mL (0.5 mM Zn^2+^, *ρ*_s,0.5mM_=0.848 ± 0.008; p=0.0008) and to 0.05 μg/mL (1.0 mM Zn^2+^,*ρ*_s,1.0mM_=0.422 ± 0.150; p=0.0218) (Figure 2).

**Fig. 2:**
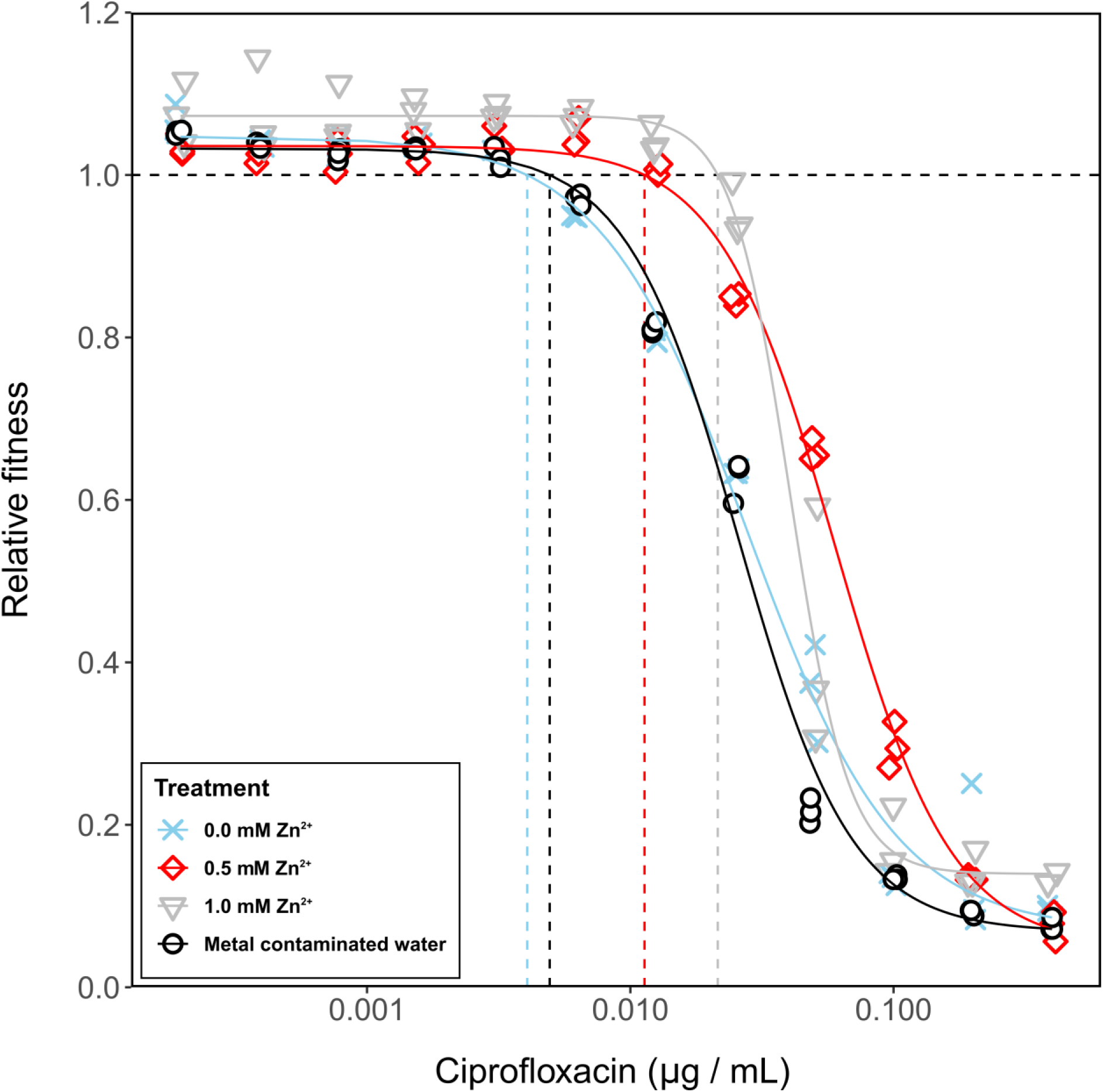
Fitness of a ciprofloxacin susceptible *E. coli* strain. relative to fitness of an isogenic ciprofloxacin resistant strain across a gradient of ciprofloxacin for three LB broth treatments amended with Zn^2+^ and one metal contaminated water source supplemented with LB. Horizontal dashed line indicates different strains perform equally well (*ρ*_s_ = *ρ*_r_ = 1), and the vertical lines represent the corresponding minimal selective concentration for the different Zn treatments.

We fitted a log-logistic dose response model with curves fitted for different Zn^2+^ concentrations (Figure 2). This significantly improved the fit compared to a reduced model (F_51,13_, p<0.001), suggesting that increasing Zn^2+^ concentrations alter the dose response relationship. Based on the full model, MSCs were estimated as ρ_s_=1 (no selection) and significantly increased from 0.0041 ± 0.0005 μg/mL in the absence of Zn^2+^ to 0.0113 ± 0.0013 μg/mL (0.5 mM Zn^2+^) and further to 0.0215 ± 0.0014 μg/mL (1.0 mM Zn^2+^) (Table 1). Zn^2+^ increased the MSC of the ciprofloxacin-susceptible strain, but also affected relative growth rate of the ciprofloxacin-resistant strain: while at 0.2 and 0.4 μg/mL ciprofloxacin growth significantly decreased (p=0.002-0.0498, df=4, *t*-test) for no Zn^2+^ and 0.5 mM of Zn^2+^, no such decrease was detected when exposed to 1 mM of Zn^2+^ (p=0.114-0.194, df=4, *t*-test).

**Table 1:** Minimal selective concentrations (MSC) of ciprofloxacin. modelled for the different Zn^2+^ concentrations and metal contaminated water

| Medium | MSC | Lower 95% CI | Upper 95% CI |
| --- | --- | --- | --- |
| **0.0 mM $\text{Zn}^{2+}$** | $0.0041 \pm 0.0005 \text{ } \mu\text{g/mL}$ | 0.0030 | 0.0051 |
| **0.5 mM $\text{Zn}^{2+}$** | $0.0113 \pm 0.0013 \text{ } \mu\text{g/mL}$ | 0.0087 | 0.0139 |
| **1.0 mM $\text{Zn}^{2+}$** | $0.0215 \pm 0.0014 \text{ } \mu\text{g/mL}$ | 0.0188 | 0.0242 |
| **Metal contaminated water** | $0.0049 \pm 0.0006 \text{ } \mu\text{g/mL}$ | 0.0037 | 0.0062 |

### Effect of Zinc on selection for gentamicin resistance

To test if the impact of Zn^2+^ on selection for antibiotic resistance is specific to ciprofloxacin or acting more generally across antibiotics, we repeated the experiment described above with gentamicin as the focal antibiotic. Isogenic *E. coli* strains with and without gentamicin resistance were grown in isolation across a gradient of gentamicin and Zn^2+^ concentrations for 24h. Growth of the gentamicin-resistant strain was reduced to 90.3 ± 1.3% relative to its susceptible counterpart in the absence of antibiotics and metals (p=0.0001, two-tailed *t*-test against 1). This cost of resistance was also apparent in the presence of Zn^2+^ (0.5mM: 90.7 ± 1.0%, p=0.0001; 1.0 mM: 90.4 ± 3.6%, p=0.0005) (Figure 3). Increasing Zn^2+^ concentrations significantly decreased growth of both the resistant and sensitive strains in the absence of gentamicin (18.7 ± 1.0% at 0.5 mM, p<0.0001; 46.6 ± 1.4% at 1 mM, p<0.0001). However, unlike ciprofloxacin, the dose response curves estimated for the different Zn^2+^ concentrations were not significantly different (model compared to reduced model: F_8,104_ =0.90, p=0.52; Figure 4).

**Fig. 3:**
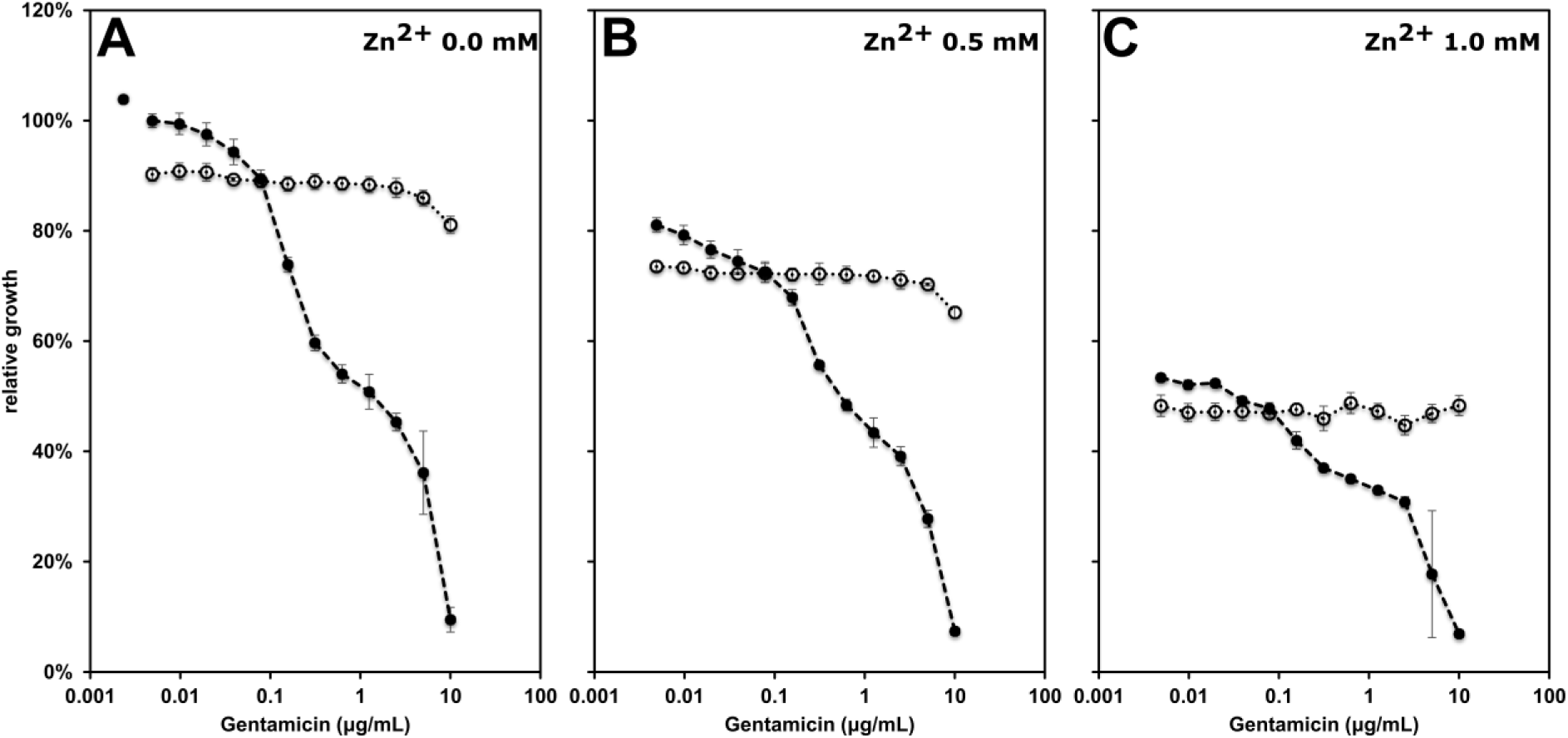
Relative growth of the susceptible (filled circles) and the gentamicin resistant (open circles) strain across a gradient of gentamicin and zinc concentrations. Growth was quantified based on OD_600_ after 24 h of growth at 37°C (n=3, average ± SD) compared to the susceptible strain in the absence of antibiotic or metal.

**Fig. 4:**
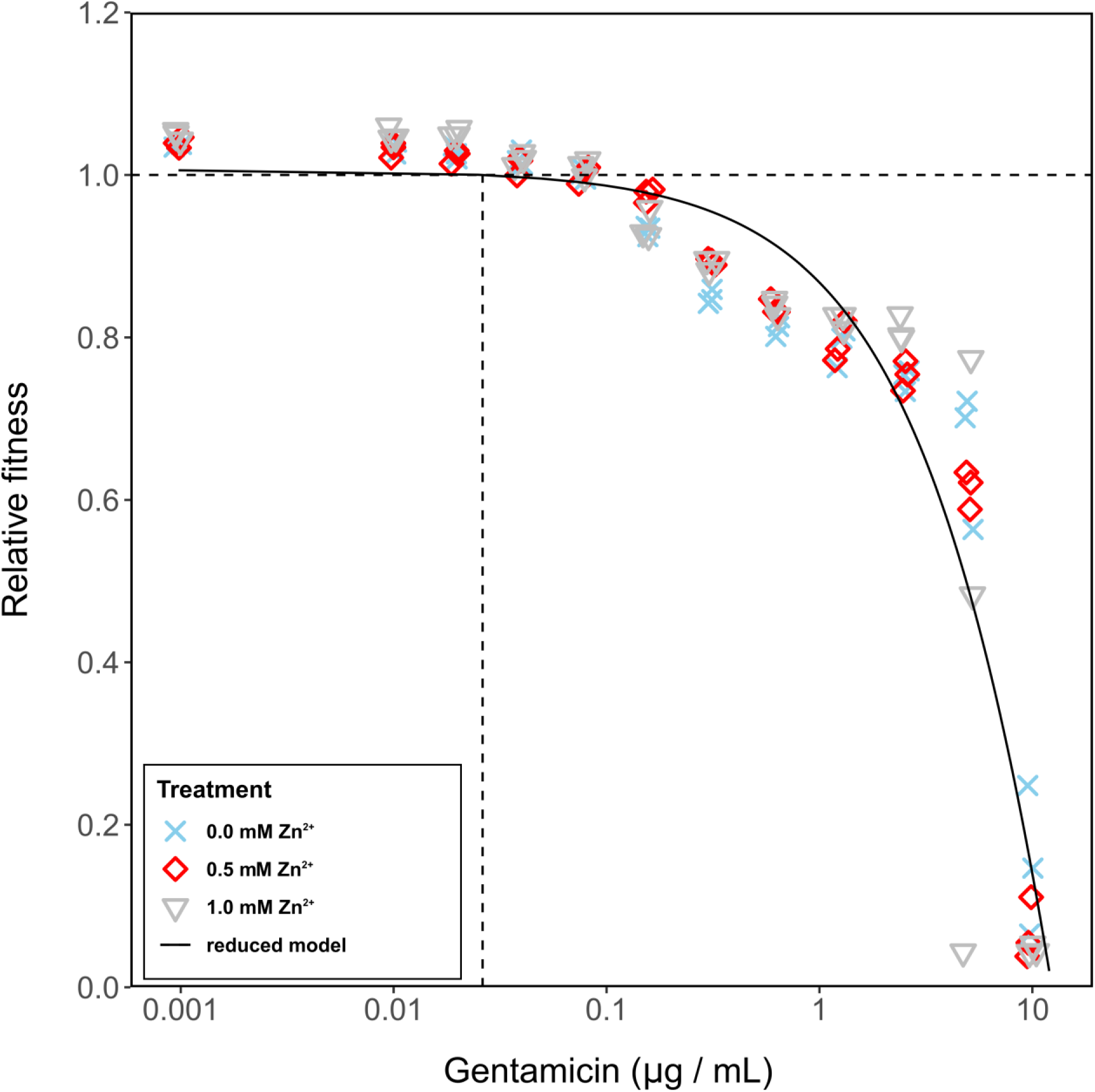
Relative fitness of the gentamicin susceptible strain across a gradient of gentamicin and zinc concentrations. Horizontal dashed line indicates no selection at a relative fitness of *ρ*_s_ = *ρ*_r_ = 1. The intercept with the reduced log-logistic dose response model indicates the minimal selective concentration illustrated with a vertical dashed line. Since no significant influence of Zn^2+^ on the log-logistic dose response model was detected exclusively the reduced models selection curve with MSC for gentamicin at 0.0242 ± 0.0162 μg/mL is shown.

### Effect of environmental metal contamination on selection for ciprofloxacin resistance

To explore the relevance of these results in more natural settings, we performed a competition experiment for ciprofloxacin using sterilized water from a tributary of the metal-contaminated Carnon River (Cornwall, United Kingdom) amended with LB. The surrounding area has a long history of mining-related heavy metal contamination (Pirrie *et al*. 2003; Environment Agency UK 2019), resulting in contamination of soil and water with a complex mixture of metals, including zinc (Table 2). Compared to LB medium made up from deionized water, the contaminated water significantly decreased relative growth of both the ciprofloxacin-resistant (77.9 ± 1.0%, p=0.0007, two-tailed *t*-test against 1) and ciprofloxacin-sensitive strain (74.2 ± 0.6% p=0.0002) with the sensitive strain having a greater relative fitness in the absence of ciprofloxacin (*ρ*_s_=1.052 ± 0.003; p=0.0012, two-tailed *t*-test against 1). Growth inhibition was quantitatively and qualitatively similar to that observed for 0.5 mM Zn^2+^ (21.3 ± 4.1% p=0.43, DF=10, *t*-test) (Figure 1D). However, the minimal selective concentration estimated from the dose response curve model was at 0.0049 ± 0.0006 μg/mL ciprofloxacin, similar to that observed for broth without any Zn^2+^ amendments (Table 2). A significant decrease in growth rate of the resistant strain at high concentrations of 0.4 μg/mL ciprofloxacin was observed (p=0.0094, *t*-test) (Figure 1).

**Table 2:** Metal concentrations in metal contaminated water from the Carnon river near Bissoe, Cornwall, UK measured by ICP-MS

| Metal | Concentration | Molarity |
| --- | --- | --- |
| **Mn** | $443.20 \pm 11.99 \text{ ng/mL}$ | $8.058 \pm 0.218 \text{ } \mu\text{M}$ |
| **Fe** | $2.50 \pm 0.43 \text{ ng/mL}$ | $0.045 \pm 0.008 \text{ } \mu\text{M}$ |
| **Ni** | $60.44 \pm 3.90 \text{ ng/mL}$ | $1.007 \pm 0.065 \text{ } \mu\text{M}$ |
| **Cu** | $439.19 \pm 3.98 \text{ ng/mL}$ | $6.971 \pm 0.063 \text{ } \mu\text{M}$ |
| **Zn** | $1837.40 \pm 21.12 \text{ ng/mL}$ | $27.839 \pm 0.320 \text{ } \mu\text{M}$ |
| **As** | $0.32 \pm 0.02 \text{ ng/mL}$ | $0.004 \pm 0.000 \text{ } \mu\text{M}$ |
| **Cd** | $2.52 \pm 0.14 \text{ ng/mL}$ | $0.023 \pm 0.001 \text{ } \mu\text{M}$ |
| **Pb** | $0.05 \pm 0.01 \text{ ng/mL}$ | $0.000 \pm 0.000 \text{ } \mu\text{M}$ |

To confirm results from the growth rate assays, strains were competed for seven days across four ciprofloxacin concentrations in the presence and absence of 0.5mM Zn^2+^ and in metal contaminated water (Figure 5). Again, relative fitness did not significantly differ between the unpolluted control and metal contaminated water at any of the tested ciprofloxacin concentrations (all p>0.05, t-test). While metal contaminated water had an effect on the growth rates of both sensitive and resistant strains, no effect on selection for ciprofloxacin resistance was observed in the environmental sample. This is most likely explained by the fact that the total metal content was high enough to affect growth, while the concentration of Zn^2+^ (0.028 mM; Table 1) was too low to significantly affect the action of ciprofloxacin. The seven-day competition experiment supported the results of the 24-hour growth rate experiment in that it showed the competitive advantage of a ciprofloxacin-susceptible strain in the presence of Zn^2+^. This is evidenced by the fact that at 0.5 mM Zn^2+^, the susceptible strain significantly outcompetes the resistant strain at 0.025 μg/mL ciprofloxacin (*ρ*_s,0.5mM_=1.084 ± 0.017), whereas in the absence of Zn^2+^ the resistant strain is more competitive (*ρ*_s,0.0mM_=0.863 ± 0.069; p=0.0059, df=4, *t*-test) (Figure 5). The effect of Zn^2+^ is apparent at concentrations as low as 0.01 μg/mL ciprofloxacin (*ρ*_s,0.5mM_=1.177 ± 0.078; *ρ*_s,0.0mM_=1.052 ± 0.014; p=0.0526, df=4) (Figure 5).

**Fig. 5:**
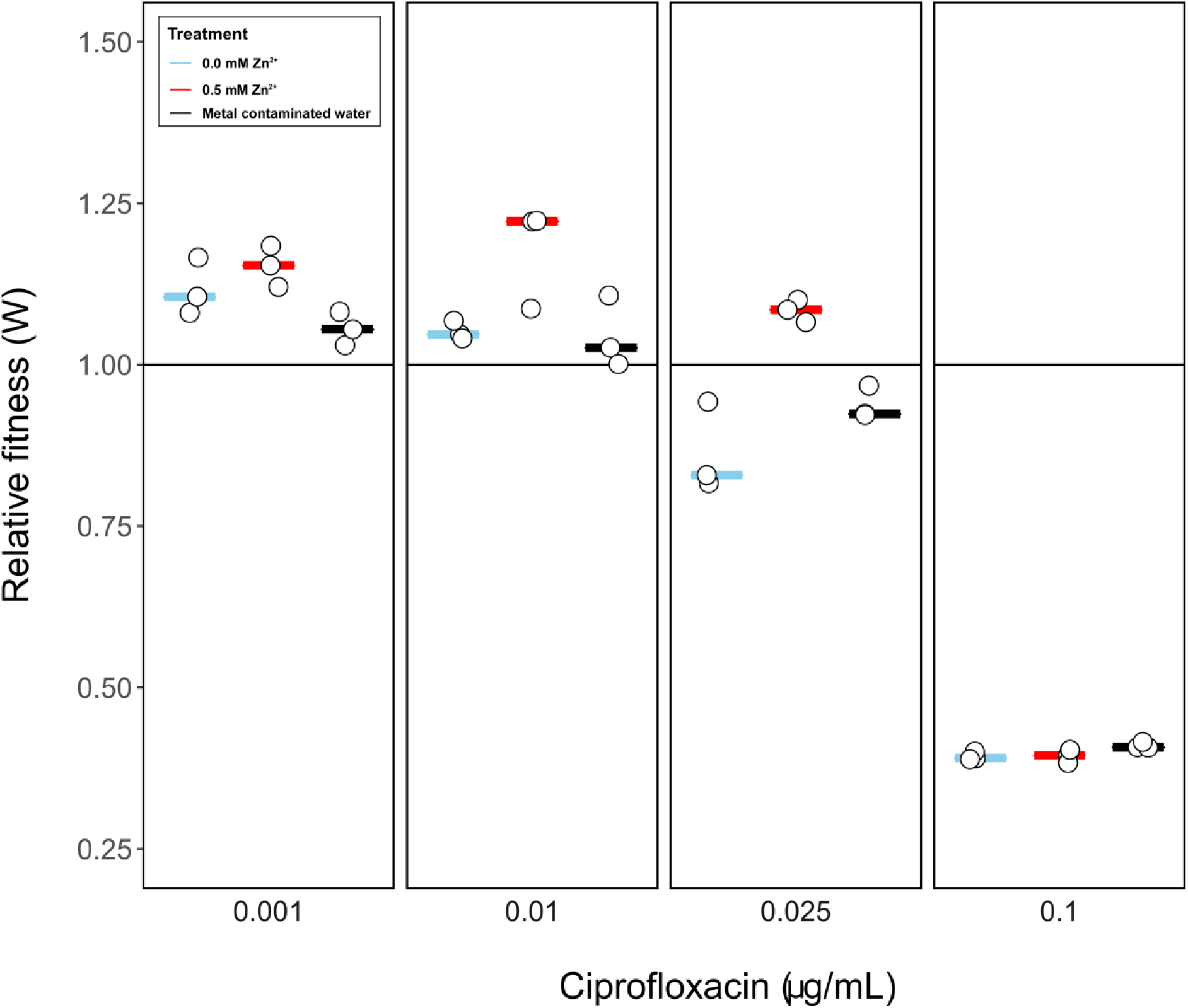
Relative fitness of the ciprofloxacin-susceptible strain based on 7-day competition experiments across a gradient of ciprofloxacin in the absence and presence of 0.5 mM Zn^2+^ and in naturally metal contaminated water.

## Discussion

Here, we present evidence that metal cations have the potential to reduce selection for ciprofloxacin resistance. The interaction of ciprofloxacin and Zn^2+^ cations, likely via chelation, resulted in a five-fold increase of the MSC for ciprofloxacin resistance at a concentration of 1.0 mM Zn^2+^. Further, metal-imposed growth inhibition decreased total population sizes of both strains. Hence, the ciprofloxacin-resistant strains exhibited decreased persistence, which resulted in an additional decrease in total incidence of resistant phenotypes. The non-fluoroquinolone antibiotic gentamicin, which does not chelate metal, did not exhibit a similar interaction with Zn^2+^, which is consistent with a scenario where zinc reduces selection for ciprofloxacin resistance by making this antibiotic unavailable through chelation.

The effect of Zn^2+^ on selection for ciprofloxacin resistance was apparent at relatively high metal concentrations (0.5 - 1 mM) in artificial media. Experiments performed using a broth-supplemented metal-contaminated water source did not reveal an effect on selection dynamics. This can be explained by the fact that the environmental Zn^2+^ concentration was over an order of magnitude lower (0.028 mM, Table 1) than the lowest defined concentration used (even though it co-occurred with a range of other metals implicated in fluoroquinolone chelation (Ma, Chiu and Li 1997)). It is possible other more heavily polluted sites have metal concentrations high enough to interfere with selection for resistance to low levels of fluoroquinolone. For example, landfill leachates have been shown to contain Zn^2+^ concentrations reaching up to 3.8 mM (Roy 1994), which we have demonstrated is within the range of concentrations affecting selection for ciprofloxacin resistance.

Our results could have important implications for human or veterinary medicine. For instance, prevalence of ciprofloxacin resistance can reach up to 21.2% of all Enterococci isolates in pig manure samples (Hölzel *et al*. 2010), making any factors altering selection dynamics highly relevant. Zinc compounds are regularly used as agricultural feed additives and growth promoters in agriculture (Poulsen 1998; Castillo *et al*. 2008) with concentrations in liquid pig manure reaching up to 4 mmol/kg wet weight (Hölzel *et al*. 2012), considerably higher than the concentrations used here. In humans, where ciprofloxacin is prescribed for a wide range of infections caused by both gram-negative and gram-positive bacteria (Redgrave *et al*. 2014), the effects observed here could potentially be relevant when patients take zinc supplements, for which daily doses of as high as 40 mg have been reported (Liu *et al*. 2017).

The focus of this study was exclusively on selection for pre-existing antibiotic resistance genes. Co-occurrence of metal and antibiotic stressors could, however, have additional effects on *de novo* evolution of antibiotic resistance. Metal stress could for example increase the rate at which antibiotic and metal resistance evolves through increased mutation rate (Lemire, Harrison and Turner 2013), which could subsequently be fixed in the population through antibiotic or metal selection. Increased spread of resistance could also result from metals impacting plasmid transfer (Klümper *et al*. 2017, 2019a).

In summary, we have shown that a selective window exists in which metals could reduce the strength of selection operating on ciprofloxacin resistance. It remains to be tested whether these conditions occur in real-world pollution scenarios. Our data highlight that in order to assess the risks of environmental AMR evolution associated with pollution with both metals and antibiotic residues it will be necessary to take complex mixtures of antimicrobial compounds into account (Ye *et al*. 2017). It is highly likely that additional metal-antibiotic interactions affect the efficacy of antibiotics (Li, Nix and Schentag 1994), and hence deserve future research attention.

## Acknowledgments

The authors thank Mr Louis Jolly for support during laboratory experiments.

